# Engineering luminopsins with improved coupling efficiencies

**DOI:** 10.1101/2023.11.22.568342

**Authors:** Ashley Slaviero, Nipun Gorantla, Jacob Simkins, Emmanuel L Crespo, Ebenezer C Ikefuama, Maya O Tree, Mansi Prakash, Andreas Björefeldt, Lauren M Barnett, Gerard G Lambert, Diane Lipscombe, Christopher I Moore, Nathan C Shaner, Ute Hochgeschwender

## Abstract

**Significance:** Luminopsins (LMOs) are bioluminescent-optogenetic tools with a luciferase fused to an opsin that allow bimodal control of neurons by providing both optogenetic and chemogenetic access. Determining which design features contribute to the efficacy of LMOs will be beneficial for further improving LMOs for use in research.

**Aim:** We investigated the relative impact of luciferase brightness, opsin sensitivity, pairing of emission and absorption wavelength, and arrangement of moieties on the function of LMOs.

**Approach:** We quantified efficacy of LMOs through whole cell patch clamp recordings in HEK293 cells by determining coupling efficiency, the percentage of maximum LED induced photocurrent achieved with bioluminescent activation of an opsin. We confirmed key results by multielectrode array (MEAs) recordings in primary neurons.

**Results:** Luciferase brightness and opsin sensitivity had the most impact on the efficacy of LMOs, and N-terminal fusions of luciferases to opsins performed better than C-terminal and multi-terminal fusions. Precise paring of luciferase emission and opsin absorption spectra appeared to be less critical.

**Conclusions:** Whole cell patch clamp recordings allowed us to quantify the impact of different characteristics of LMOs on their function. Our results suggest that coupling brighter bioluminescent sources to more sensitive opsins will improve LMO function. As bioluminescent activation of opsins is most likely based on Förster resonance energy transfer (FRET), the most effective strategy for improving LMOs further will be molecular evolution of luciferase-fluorescent protein-opsin fusions.

## 1 Introduction

Manipulation of neuronal activity in behaving experimental animals is crucial for elucidating neuronal networks underlying brain function. Both optogenetic^1^ and chemogenetic^2^ approaches continue to be highly valuable for identifying the contributions of genetically-defined neuron populations to circuit and behavioral outputs. Both methods offer distinct advantages, with optogenetic control of the activity of subsets of neurons at precise temporal scales and slower chemogenetic control of the activity of entire populations of neurons across the brain. Previous work has developed a toolset that integrates opto- and chemogenetic approaches by fusing a light-emitting luciferase to an optogenetic light-responsive element, resulting in a luminescent opsin, or luminopsin (LMO)^3-5^ [Fig. 1(a)]. Bioluminescence produced by oxidation of a diffusible luciferin substrate by the luciferase enzyme activates a tethered, nearby opsin. Depending on the biophysical properties of the opsin, the light generated by the luciferase can either excite or inhibit target neurons expressing the LMO. Such integration of opto- and chemogenetic approaches allows manipulation of neural activity over a range of spatial and temporal scales of the same neurons in the same experimental animal. For example, the contribution to behavioral components of the activation of an entire population of neurons can be compared to those of a subset of the same neurons, activating the opsin chemogenetically through bioluminescence or optogenetically through light fibers^6^.

**Fig. 1.**
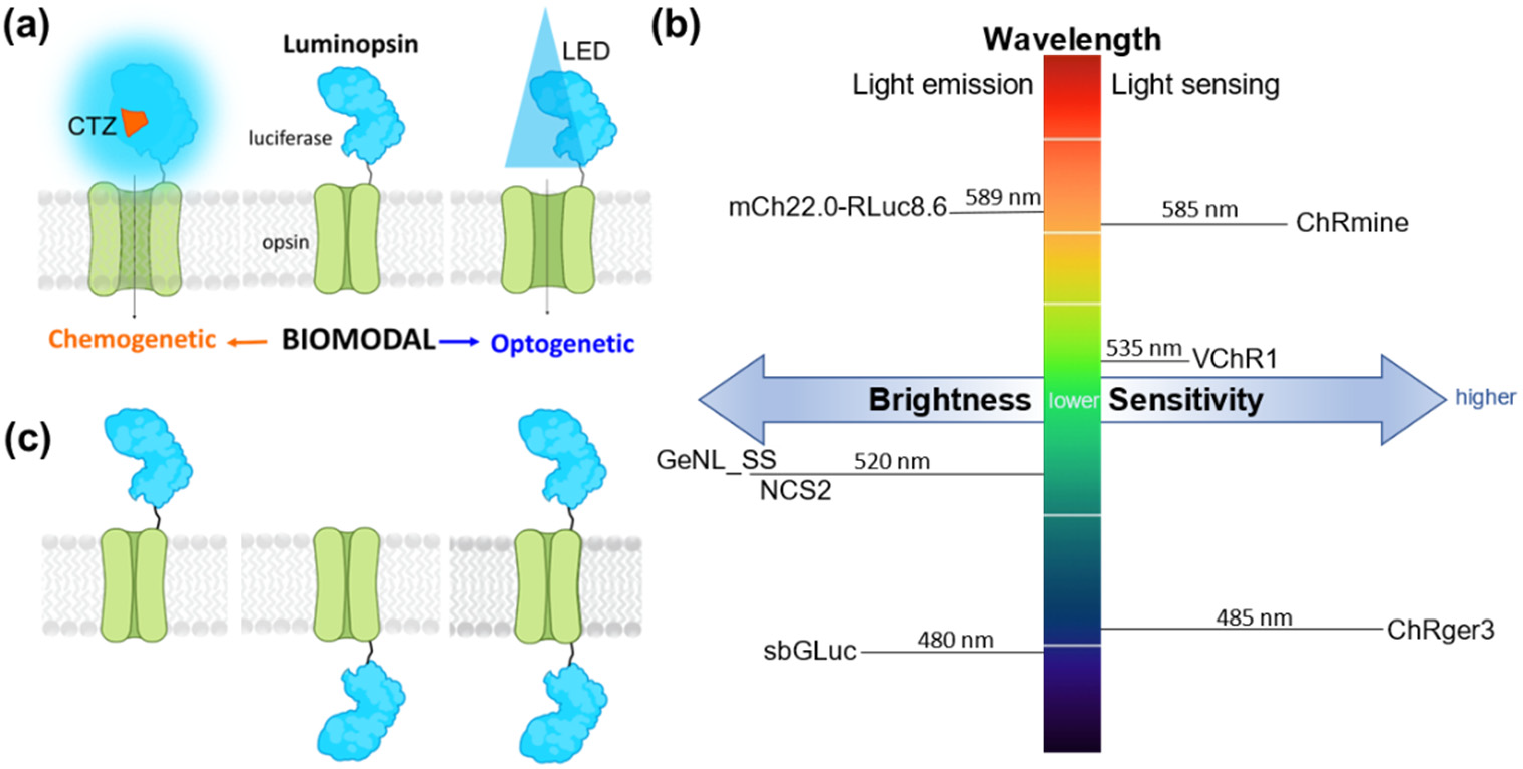
Luminopsin (LMO) features. (a) The luciferase-opsin fusion molecule (luminopsin) enables bimodal chemogenetic (CTZ) and optogenetic (LED) access to opsin activation. (b) Parameters for improving LMO performance are brightness of the light emitter, light sensitivity of the light sensor, and matching spectra for peak light emission and sensing. Moieties used in this study are indicated. (c) Possibilities for arrangement of moieties are N-terminal, C-terminal, or N- and C-terminal fusion of the luciferase to the opsin.

For the first excitatory LMO1, wild type luciferase (GLuc)^7^ was combined with channelrhodopsin 2 (ChR2)^8^. While application of luciferin induced small inward currents in HEK293 cells, bioluminescence was not able to generate action potentials in neurons^3^. Efficiently changing membrane potentials in neurons required the use of a brighter light emitter such as the GLuc variant ‘slow burn’ (sbGLuc)^9^ in combination with optogenetic channels with higher light sensitivity, such as *Volvox* channelrhodopsin 1 (VChR1)^10-11^. This combination in LMO3 yielded robust and reliable activation of neurons in numerous *in vivo* experiments^5-6,12-19^. We and others have since continued to combine more potent light emitters with more light sensitive opsins to generate LMOs with still higher efficacies. These efforts include use of novel luciferase mutants, such as GLucM23^20^, molecularly evolved luciferase-fluorescent protein fusions, such as NCS2 (eKL9h-mNeonGreen)^21^, and opsins with higher light sensitivities, such as step function opsins^22^. These further improvements in LMO efficacy enable more effective neuron activation with less luciferin or via easier application routes (for example, intraperitoneal instead of intravenous).

Taking advantage of new developments and discoveries in optogenetic tools, such as the machine learning-guided engineering of the designer channelrhodopsin ChRger^23^ and the discovery of the marine opsin ChRmine from *Tiarina fusus*^24^ together with recent progress in further evolving luciferases (SSLuc^25^), we have continued to generate new versions of LMOs (Table 1). Design principles in addition to brightness of luciferases and sensitivity of opsins were the arrangement of moieties and the wavelength compatibility of emitter and sensor [Fig. 1 (b, c)]. The moieties of LMOs can be arranged in one of three ways, with the luciferase on the N-terminus, C-terminus or both the N- and C-termini of the opsin [Fig. 1 (c)]. In all our previous designs the luciferase was tethered to the N-terminus of the opsin. However, for the inhibitory iLMO2^4^ the RLuc-based luciferase–fluorescent fusion protein Nanolantern^26^ was tethered to the C-terminus of the opsin *Natronomonas* halorhodopsin (NpHR)^27^. Each luciferase and opsin partnered for an LMO has a unique peak emission and absorption spectrum, respectively [Fig. 1(b)]. We hypothesized that closely matching these spectra will improve the function of LMOs.

**Table 1.** Summary of luminopsins assessed

| Name | Light Emitter<br>(Peak Em Wavelength) | Opsin<br>(Peak Ex Wavelength) | Arc Lamp<br>(Ex) | Arrangement of<br>Light Emitter | Fluorescent<br>Protein | Luciferin |
| --- | --- | --- | --- | --- | --- | --- |
| LMO3 | sbGLuc (480 nm) | VChR1 (535 nm) | Blue (480 nm) | N-terminal | EYFP | CTZ |
| LMO7 | NCS2 (520 nm) | VChR1 (535 nm) | Green (540 nm) | N-terminal | mNeonGreen | hCTZ |
| LMO7.2 | NCS2 (520 nm) | VChR1 (535 nm) | Green (540 nm) | C-terminal | mNeonGreen | hCTZ |
| LMO7.3 | NCS2 (520 nm) | VChR1 (535 nm) | Green (540 nm) | N- and C-terminal | mNeonGreen | hCTZ |
| LMO8 | NCS2 (520 nm) | ChRger3 (485 nm) | Green (540 nm) | N-terminal | mNeonGreen | hCTZ |
| LMO10 | GeNL_SS (520 nm) | VChR1 (535 nm) | Green (540 nm) | N-terminal | mNeonGreen | hCTZ |
| LMO11 | GeNL_SS (520 nm) | ChRmine (585 nm) | Green (540 nm) | N-terminal | mNeonGreen | hCTZ |
| LMO12 | mCh22.0_RLuc8.6 (589 nm) | ChRmine (585 nm) | Red (580-630 nm) | C-terminal | mCh22.0 | eCTZ |

To compare the efficacy of different LMOs in changing a cell’s membrane potential, we assessed eight excitatory LMOs through whole cell patch clamp recordings in HEK293 cells. By measuring the amplitude of inward current elicited with a metal halide light source versus the one elicited by bioluminescence, we were able to calculate for each LMO the coupling efficiency, the percentage of optogenetic activation of an opsin that can be achieved with bioluminescent stimulation^3^. We used this quantification of the efficacy of a light emitter-light sensor pairing to identify which design characteristics of LMOs are most impactful.

## 2 Materials and Methods

### 2.1 Plasmids

Plasmids were cloned with conventional molecular biology techniques. Some constructs were cloned into both pcDNA and pAAV vectors for testing in HEK293 cells and primary neurons, respectively. Constructs are listed in Table 2.

**Table 2.** Plasmids used in this paper

| pcDNA Constructs | pAAV Constructs | Addgene number |
| --- | --- | --- |
| pcDNA3.1-CAG-LMO3 (sbGLuc-VChR1-EYFP) | pAAV-hSyn-LMO3 (sbGLuc-VChR1-EYFP) | 114099 (pAAV) |
| pcDNA3.1-CAG-LMO7 (mNG-eKL9h-VChR1) | pAAV-hSyn-LMO7 (mNG-eKL9h-VChR1) | 205097 (pAAV) |
| pcDNA3.1-CAG-LMO7.2 (VChR1-mNG-eKL9h) |  |  |
| pcDNA3.1-CAG-LMO7.3 (mNG-eKL9h -VChR1-mNG-eKL9h) |  |  |
| pcDNA3.1-CAG-LMO8 (mNG-eKL9h-ChRger3) |  |  |
| pcDNA3.1-CMV-LMO10 (GeNL_SS-VChR1) |  |  |
| pcDNA3.1-CAG-LMO11 (GeNL_SS- ChRmine) | pAAV-hSyn-LMO11 (GeNL_SS-ChRmine) | 213034 (pAAV) |
| pcDNA3.1-CAG-LMO12 (ChRmine-mCh22.0-RLuc8.6) |  |  |

### 2.2 Virus

AAVs carrying LMOs were produced by transfecting subconfluent HEK293FT cells per 10-cm culture dish with 24 μg of the helper plasmid pAd delta F6, 20 μg of the serotype plasmid AAV2/9, and 12 μg of the LMO plasmid using Lipofectamine 2000. After 72 h, the supernatant was harvested from culture plates and filtered at 0.45 μm. Virus was purified from cells and supernatant according to previously described methodology^28^, but without the partitioning step in the aqueous two-phase system. Virus was dialyzed against PBS (w/o Ca, Mg) overnight at 4 °C, using FLOAT-A-LYZER G2, MWCO 50 kDa, followed by concentration in Amicon Ultra-0.5 379 mL Centrifugal Filters. Viral titers were determined by quantitative PCR for the woodchuck hepatitis post-transcriptional regulatory element. Preparations with titers around 1 × 10^13^ vg/mL were used in the study.

### 2.3 Cell Culture

#### 2.3.1 HEK293 cells

Human embryonic kidney fibroblasts (HEK293, RRID:CVCL_0045) were grown in Dulbecco’s Modified Eagle Medium (DMEM, ThermoFisher) supplemented with 10% fetal calf serum (FBS, ThermoFisher) and 0.5% penicillin/streptomycin (ThermoFisher). Cells were cultured in T25 flasks with gas exchange caps at 37°C and 5% CO_2_ atmosphere. Cells were split once a week by trypsinization (0.25%, Gibco, ThermoFisher). Cells were seeded at full confluency on a 6-well plate. Transfection was done by lipofection (Lipofectamine 2000, ThermoFisher) with 2 μg of plasmid for 4-6 hours. Cells were then trypsinized (TrypLE Express, Gibco, ThermoFisher) and seeded across six wells of a 12-well plate on 15 mm PDL coated coverslips (Neuvitro).

#### 2.3.2 Primary neurons

Rat embryonic day 18 primary cortical neurons were recovered from tissue samples as per vendor’s instructions (TransnetYX) and plated in Neurobasal Medium (Invitrogen) containing 2% B-27 (Invitrogen), 2 mM Glutamax (Invitrogen), 5% FCS, and 0.5% gentamycin (NB-FCS). Cells were plated on 1-well MEA plates (60MEA200/30iR-Ti; Multichannel Systems) at a seeding density of 7x10^4^ in a 10 μl drop^29,30^. Once the cells had adhered to the surface, the wells were slowly flooded with NB-FCS and plates were returned to the incubator. Viral transduction of constructs occurred the next day (∼18-24h) together with changing to culture medium without serum (NB-Plain: Neurobasal Medium, 2% B-27, 2 mM Glutamax, 0.1% gentamycin). Three days later media was changed to Neurobasal Plus Medium (Invitrogen), 2% B-27 Plus supplement (Gibco, ThermoFisher), 2 mM Glutamax, 0.1% gentamycin (NB+). Every 3-4 days a third of media was changed out with fresh NB+. Cells were maintained in a 5% CO_2_ atmosphere at 37°C.

### 2.4 HEK Cell Electrophysiology

Whole cell voltage patch clamp was performed on HEK293 cells 24-72 hours after transfection. A coverslip was transferred to a recording chamber (RC26-GLP, Warner Instruments) on the stage of an upright microscope (BX51WI, Olympus) and perfused with external buffer (1.5 mL/min) containing (in mM): 150 NaCl, 3 KCl, 10 HEPES, 2 CaCl_2_, 2 MgCl_2_ and 20 D-glucose (pH 7.4, ∼300-315 mOsm/kg) at a temperature of 37°C ± 1°C. Intracellular patch solution contained (in mM): 130 K-gluconate, 8 KCl, 15 HEPES, 5 Na_2_-phosphocreatine, 4 Na_2_-ATP, 0.3 Na_2_-GTP (pH 7.25, ∼295-300 mOsm/kg). Borosilicate glass micropipettes were prepared using a vertical pipette puller (PC-100, Narishige) and had resistances from 2-6 MΩ. Epifluorescence microscopy was used to identify cells expressing the construct (constructs contained red or green fluorescent proteins). Opsins were excited through the objective (LUMPLFLN40XW, 433 NA 0.8, Olympus) using a metal halide light source (130 W, U-HGLGPS, Olympus) with filter cubes for blue (Ex/Em: 480/530 nm, U-MNIBA3, Olympus), green (Ex/Em: 540/600, U-MWIGA3, Olympus) and red (Ex/Em: 580-630/645-695, ET-Alexa633 Filter Set 605/50x, 670/50M w/BX2 cube) excitation. An electronic shutter (Lambda SC, Sutter Instruments) was used to control light delivery to cells and limit exposure to 1s. Photocurrents were evoked with blue, green, or red light to match the peak bioluminescence emission of the luciferase tethered to the opsin. At maximum light intensity from the light source irradiance was (in mW/cm^2^) 16.8, 36.5, and 39.9, respectively, as measured with a light meter (ThorLabs).

Recordings were conducted at -60 mV holding voltage using a Multiclamp 700b amplifier and Digidata 1440 digitizer with pClamp 10 recording software (Molecular Devices). After sealing the cells, photocurrent response was measured using a gap free protocol followed by addition of luciferin via perfusion. Luciferins included h-coelenterazine (hCTZ, Nanolight Technology, #301), native-coelenterazine (CTZ, Nanolight Technology, #303), and e-coelenterazine (eCTZ, Nanolight Technology, #355). Luciferin was diluted to a 500 μM working stock in ACSF (500 μL total) for CTZ, hCTZ, and eCTZ. Final concentration in perfusion chamber was ∼100 μM. Luciferin was chosen based on the most efficient substrate for the respective luciferase in the tested construct.

### 2.5 Multi Electrode Array Recordings

MEA2100-Lite-System (Multichannel Systems, Germany) was used for all MEA recordings with Multichannel Experimenter software at a sample rate of 10,000 Hz. Consistently spiking neurons were used for recordings between DIVs 14–25; only cultures showing spontaneous electrophysiological activity were used. All-trans retinal (R2500; Sigma-Aldrich, St. Louis, MO) was added to the culture medium to 1 μM final concentration before electrophysiological recordings. Prior to recording, all reagents were pre-warmed to 37°C. MEAs were transferred from the CO2 incubator to the heated MEA2100 head stage maintained at 37°C, and the cultures were allowed to equilibrate for 5–10 min. The headstage was situated on a support stage with a PlexBright optical patch cable positioned from below to illuminate the center of the MEA dish. The patch cable was connected to a Plexon optogenetic LED fiber-coupled module (Blue, 465 nm; Green, 525 nm; Lime, 550 nm) controlled by a PlexBright LD-1 Single Channel LED Driver for light stimulation of cultures at different wavelength. A micropipette was used to add luciferin or vehicle with the reagent drop gently touching the liquid surface, creating a time-locked artifact in the recordings. After recording, the media in the wells was replaced with fresh pre-equilibrated and pre-warmed NB-Plain media, and cultures were used for another round of recording the next day.

### 2.6 Data Analysis

#### 2.6.1 HEK Cell Electrophysiology

Recordings were sampled at 10 kHz. Photocurrents were assessed in Clampfit (Molecular Devices) at steady state level during the last 100 ms of illumination to eliminate inconsistencies found before light-adaptation during peak current response. Luciferin-induced currents were measured in Clampfit where the amplitude plateaus during stimulation. Coupling efficiencies were determined by normalizing luciferin-induced current change to photocurrent change elicited by the lamp (ie. luciferin-induced current / photocurrent). In non-expressing controls, LED and luciferin impact on current was determined.

#### 2.6.2 MEA Recordings

All MEA analysis was done offline with Multichannel Analyzer software (Multichannel Systems) and NeuroExplorer (RRID: SCR_001818). Spikes were counted when the extracellular recorded signal exceeded 9 standard deviations of the baseline noise. For assessing the effects of CTZ (10μm final concentration), only electrodes displaying the expected change in spiking activity with blue light from the LED light source, i.e. opsin expressing neurons, were evaluated. Spikes were counted before luciferin or vehicle was added and immediately after addition. Spike frequency was also assessed for LED exposure. Pooled data was obtained from different electrodes (a) of the same culture, (b) from different cultures, and (c) over different DIVs.

To evaluate the significance of differences in firing rate across conditions and groups, we performed a permutation test. This non-parametric approach allowed us to compare neuronal firing rates between pre- and post-treatment conditions across LED, vehicle, and h-CTZ treatments in LMO7 and LMO11 expressing cultures. For each treatment data were grouped into ‘pre’ and ‘post’ conditions. For each iteration of the test, we shuffled the data simulating the null hypothesis that presumes no difference between the conditions. After shuffling, the dataset was split again into ‘pre’ and ‘post’ groups, maintaining their original sizes, and the mean difference between these groups was calculated. This process was repeated 30,000 times to construct a null distribution of mean differences, representing the expected variability under the null hypothesis. The actual mean difference was calculated between the pre and post conditions for each treatment and group, comparing it with the null distribution. The p-value was calculated based on how the actual observed difference in means compared to the null distribution. Given the multiple comparisons involved across treatments and groups, we applied a Bonferroni correction to adjust the threshold for significance.

## 3 Results

### 3.1 Coupling efficiency of LMOs

To evaluate the performance of LMOs, whole-cell patch clamp recordings were obtained from cultured HEK293 cells expressing these constructs. For each recording, opsins were activated by an arc lamp to obtain the maximal photocurrent and by luciferin (CTZ, hCTZ, eCTZ) to obtain the bioluminescence-induced current [Fig. 2(a)]. We began by measuring, under voltage clamp conditions (−60 mV), the photocurrents evoked by exciting the opsin actuators directly via illumination by an arc lamp, choosing a filter cube matching the absorption peak of the opsin. As shown in the example in Fig. 2(b), for a cell expressing LMO7, illumination evoked an inward photocurrent due to activation of VChR1. This current reached a peak within a few ms and exhibited a sustained plateau that persisted for as long as the excitation light was maintained (1 s in the example shown in Fig. 2(b)). To determine the effectiveness of the LMO for chemogenetic control of membrane potential, we measured the ability of the luciferin to induce currents in these cells. Infusion of luciferin into the recording chamber yielded bioluminescence emission and simultaneous activation of an inward current [Fig. 2(b)]. This inward current generally tracked the time course of the luciferin presence in the chamber. We then used these measurements to determine the efficiency of coupling between the luciferase light source and the opsin actuator. We employed a simplified version of the procedure introduced by Berglund et al^3^. In brief, we compared the magnitude of currents induced by luciferin to those evoked by maximal exposure to light from an arc lamp by dividing the measured luciferin-induced current by the maximal photocurrent, yielding the coupling efficiency [Fig. 2(c)]. This value defines the intrinsic efficiency of coupling between luciferase and opsin and takes into account cell-to-cell variations in LMO expression. Figure 2(d) shows examples of non-expressing control HEK293 cells patched to determine the current response to light of different wavelength and to different luciferins (CTZ, hCTZ, eCTZ). Examples are representative of 5 cells recorded for each experiment.

**Fig. 2.**
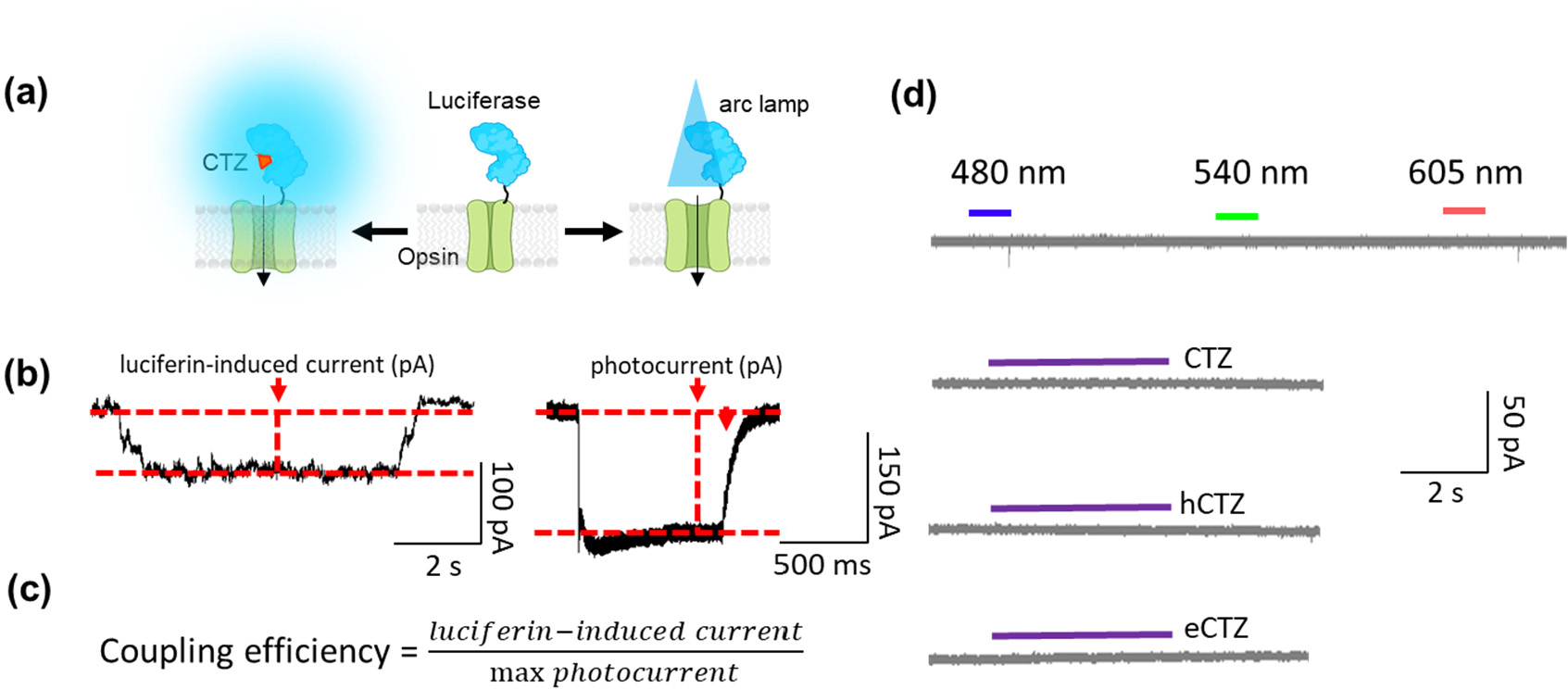
Quantifying LMO efficacy. (a) Membrane potential changes elicited chemo (CTZ)- or opto (LED, arc lamp)- genetically can be measured in HEK293 cells expressing an LMO. (b) In whole cell voltage patch clamp, inward current is determined in cells, here expressing LMO7, exposed to luciferin (hCTZ) and to light from an arc lamp (photocurrent). (c) Equation used to quantify LMO efficacy using currents measured in patch clamp. (d) Absence of effects of various luciferins and wavelengths of physical light source on the current of non-expressing HEK293 cells (controls, n=5).

### 3.2 Brightness of light emitter

In previous studies we saw a continuous improvement of LMO function when replacing a luciferase with a brighter one. A 10-fold improvement in coupling efficiency was achieved when switching out wildtye GLuc (LMO2) with the brighter variant sbGLuc (LMO3), leading to the first excitatory LMO that allowed non-invasive manipulation of animal behavior^5^. This trend continued, though not as dramatically, when we replaced sbGLuc in LMO3 with a luciferase-fluorescent protein FRET probe (NCS2, a fusion of eKL9h and mNeonGreen), resulting in LMO7^21^. Here we took advantage of a new FRET probe, GeNL_SS, a bright light emitter obtained by extensive molecular evolution of a GeNL – SSLuc fusion protein. Tethering GeNL_SS to VChR1 resulted in LMO10. Direct comparison of VChR1-based LMOs 3 (sbGLuc) and 7 (NCS2) with 10 (GeNL_SS) showed an improvement in coupling efficiency at 0.57 (n = 5) when compared to LMO7 (CE = 0.52; n = 5) and LMO3 (CE = 0.31 ; n = 5) [Fig. 3(a-d)].

**Fig. 3.**
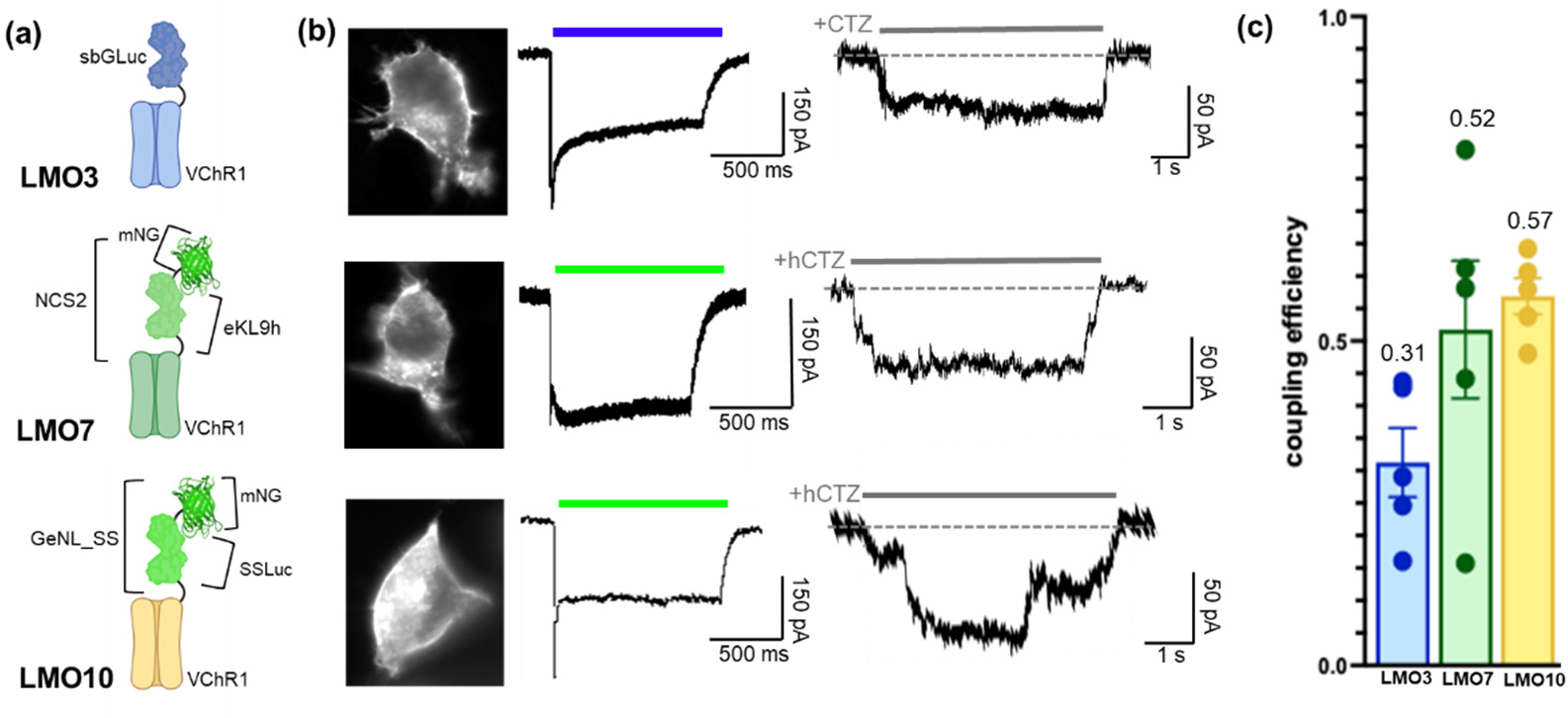
Increasing brightness of the light emitter. (a) Schematics of LMO3, LMO7, and LMO10 with same opsin (VChR1) and different light emitters (sbGLuc, NCS2, GeNL_SS). (b) Fluorescent images (left), photocurrent patch clamp traces (middle), and luciferin-induced patch clamp traces (right) for LMO3, LMO7, and LMO10. (c) Coupling efficiencies of LMO3, LMO7, and LMO10 (n=5).

### 3.3 Sensitivity of light sensor

We previously also saw an improvement of LMO function when replacing an opsin with a more light-sensitive one. Again, a 10-fold improvement in coupling efficiency was observed from LMO1 (Gluc-ChR2) to LMO2 (Gluc-VChR1)^3^. Thus, we expected to see an increase in coupling efficiency when we fused NCS2 used successfully in LMO7 (NCS2-VChR1) to the designer channelrhodopsin ChRger3^23^ in LMO8. The ChRger series of excitatory opsins was optimized through a machine learning platform yielding high-photocurrent ChRs with high light sensitivity. While ChRger3 displayed a large response to lamp stimulation, it completely lacked a response to bioluminescence [n = 5, Fig. 4(c, d)].

**Fig. 4.**
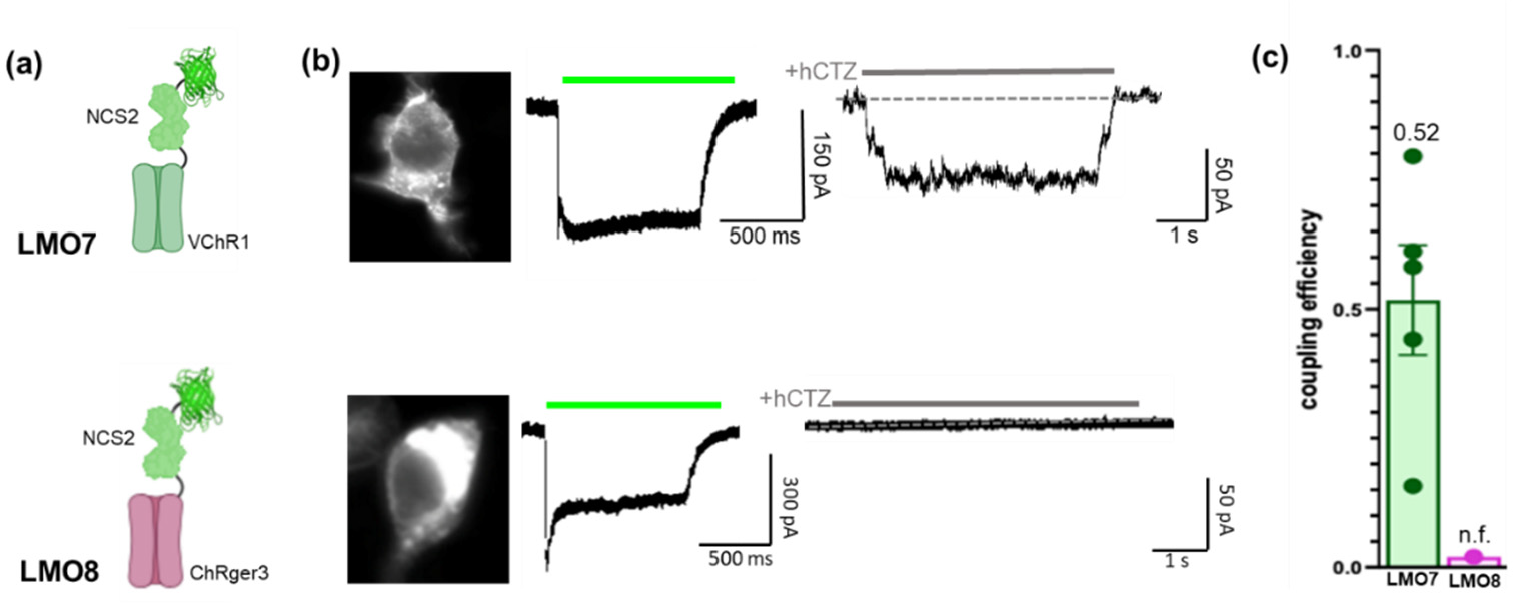
Increasing sensitivity of the light sensor. (a) Schematics of LMO7 and LMO8 with same luciferase-opsin fusion protein (NCS2) and different opsins (VChR1, ChRger3). (b) Fluorescent images (left), photocurrent patch clamp traces (middle), and luciferin-induced patch clamp traces (right) for LMO7 and LMO8. (c) Coupling efficiencies of LMO7 and LMO8 (n=5).

We next paired our currently brightest light emitter, GeNL_SS, with ChRmine, an opsin identified in a marine organism–based genomic screen for new classes of microbial opsins^24^. ChRmine was reported to have a hundred-fold improved operational light sensitivity compared with other fast, red-shifted variants. While combining GeNL_SS with VChR1 in LMO10 outperformed LMO7 (NCS2-VChR1), the combination with ChRmine in LMO11 (GeNL_SS-ChRmine) led to further improved coupling efficiency from 0.57 (LMO10) to 0.65 (LMO11) [Fig. 5 (a-d)].

**Fig. 5.**
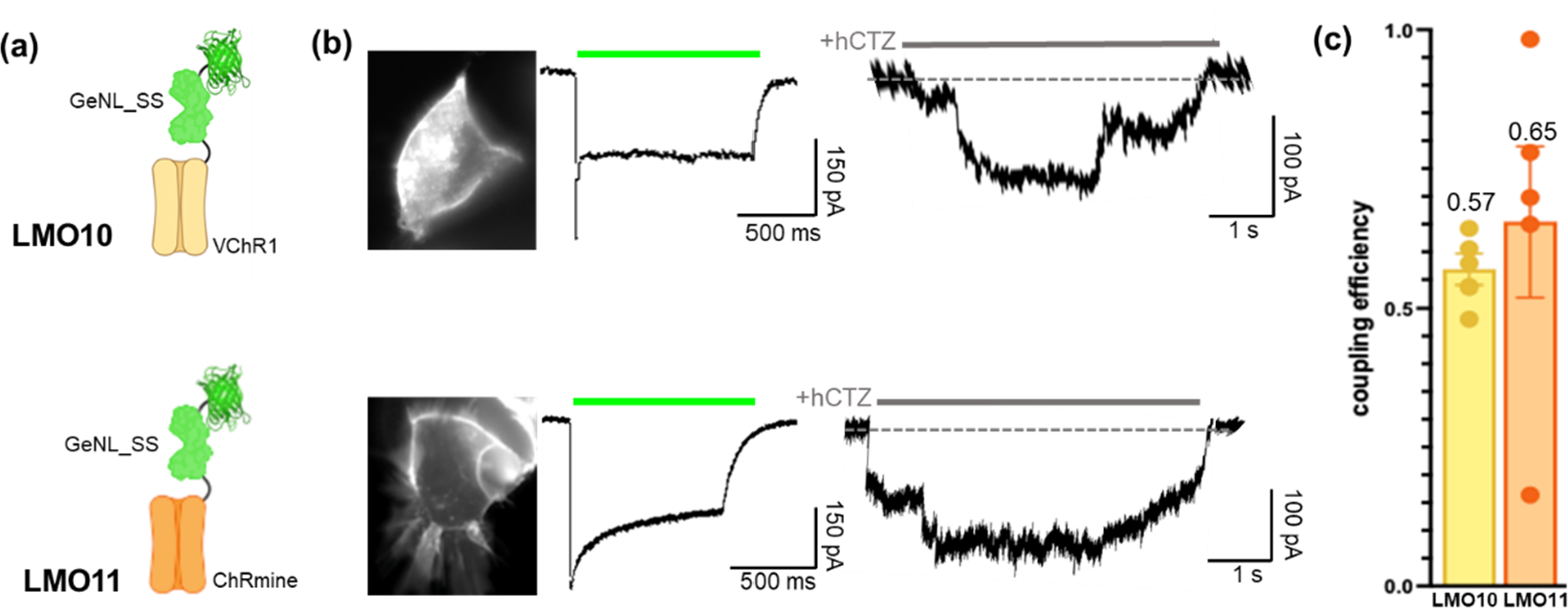
Increasing both brightness of the light emitter and sensitivity of the light sensor. (a) Schematics of LMO10 and LMO11 with brightest luciferase-opsin fusion protein (GeNL_SS) and opsins VChR1 and the super light sensitive ChRmine. (b) Fluorescent images (left), photocurrent patch clamp traces (middle), and luciferin-induced patch clamp traces (right) for LMO10 and LMO11. (d) Coupling efficiencies of LMO10 and LMO11 (n=5).

### 3.4 Arrangement of Moieties

To compare different configurations of how the light emitter is attached to the opsin we generated variants of LMO7 (NCS2-VChR1) with either NCS2 tethered to the C-terminus instead, LMO7.2 (VChR1-NCS2), or with an additional light source tethered to the C-terminus, LMO7.3 (NCS2-VChR1-NCS2) [Fig. 6(a)]. All LMO7 expressing cells showed both photocurrent and luciferin-induced current responses of variable magnitudes with a coupling efficiency of 0.56 (n = 5) [Fig. 6(c,d)]. However, the C-terminal fusion of the light emitter in LMO7.2 permits chemogenetic activation with a diminished bioluminescent response reflected in the lower coupling efficiency of 0.30 (n = 5) [Fig. 6(c,d)]. In contrast, of five LMO7.3 expressing cells, three completely lacked a response to lamp stimulation, while two had a minimal response [Fig. 6(c,d)]). None of the cells showed any luciferin-induced current. This diminished response is reflected in the nonfunctional (n.f.) coupling efficiency of LMO7.3 (n = 5).

**Fig. 6.**
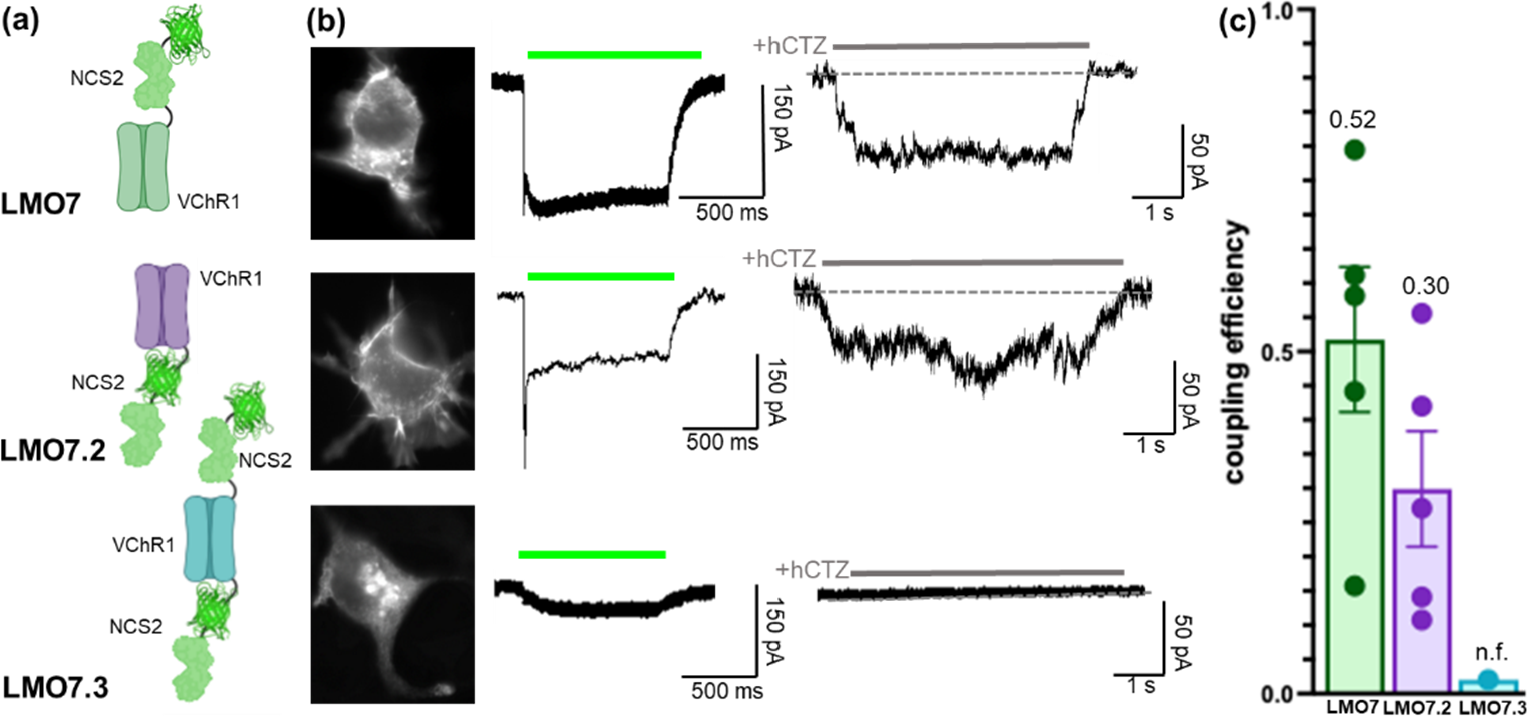
Different arrangements of moieties in LMOs. (a) Schematics of NCS2 tethered to VChR1 at the N-terminal (LMO7), the C-terminal (LMO7.2), and the N- and C-terminal (LMO7.3). (b) Fluorescent images (left), photocurrent patch clamp traces (middle), and luciferin-induced patch clamp traces (right) for LMO7, LMO7.2, and LMO7.3. (c) Coupling efficiencies of LMOs 7, 7.2, and 7.3 (n=5).

### 3.5 Wavelength compatibility

In LMO11 the peak emission of the light emitter GeNL_SS is around 520 nm, and the peak absorption of the opsin ChRmine is at 585 nm^24^. To generate an LMO with closer matched peak spectra we tethered a red-shifted light emitter with a peak emission of 589 nm, a fusion of mCherry22.0 and the red-shifted *Renilla* luciferase variant RLuc8.6(535W156F), to the C-terminus of ChRmine [Fig. 7(a)], generating LMO12. While the photocurrent generated by light exposure form the arc lamp remained the same for both LMOs, the coupling efficiency of LMO12 (CE = 0.16, n = 5) was vastly diminished compared to LMO11 (CE = 0.65, n = 5) [Fig. 7(b,c)].

**Fig. 7.**
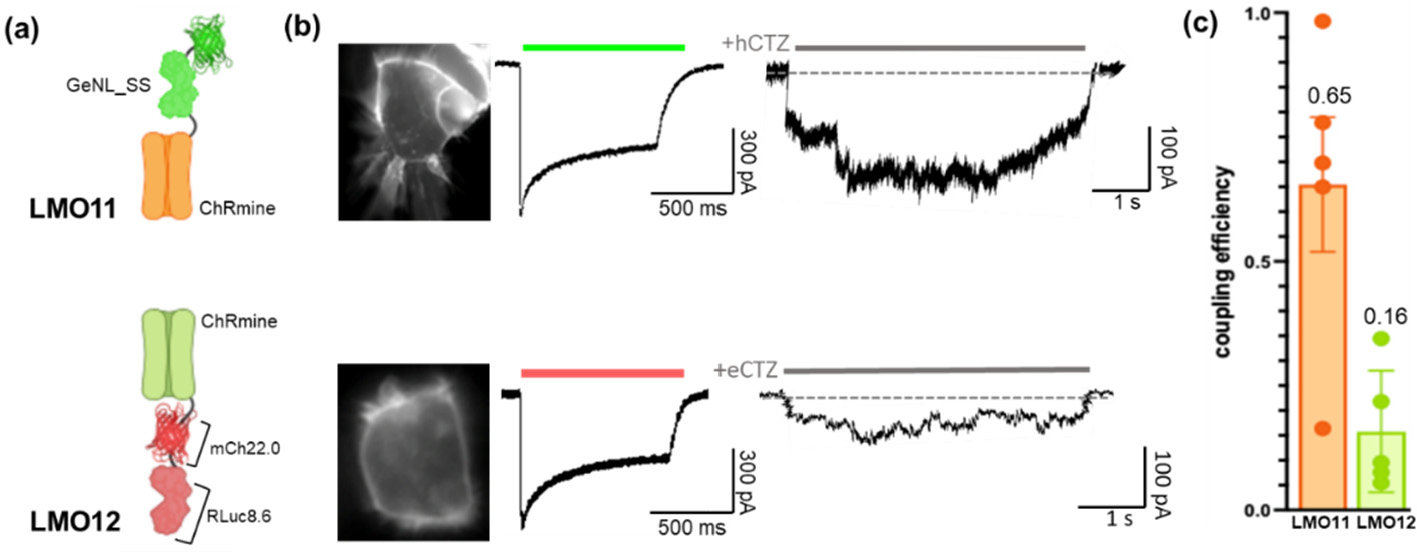
Matching emission and excitation spectra of moieties in LMOs. (a) Schematics of LMO11 (GeNL_SS-ChRmine) and LMO12 (ChRmine-mCh22.0-RLuc8.6). (b) Fluorescent images (left), photocurrent patch clamp traces (middle), and luciferin-induced patch clamp traces (right) for LMO11 and LMO12. (d) Coupling efficiencies of LMOs 11 and 12 (n=5).

### 3.6 LMO performance in activating neuronal populations

We used multi-electrode extracellular recordings to examine the ability of LMOs to activate neuronal populations. For these experiments, we transduced the entire population of rat embryonic cortical neurons in an MEA well with AAV-hSyn-LMO, using a virus titer that yielded transduction of most neurons by LMOs. Figure 8 shows the effects of LED, vehicle, and CTZ treatments in spontaneously firing cultures on neuronal firing rates in LMO7 and LMO11 expressing cortical cultures. Implementation of a permutation test, followed by a Bonferroni correction for multiple comparisons, provided a robust assessment of the significance of the observed changes between pre- and post-treatment conditions.

**Fig. 8.**
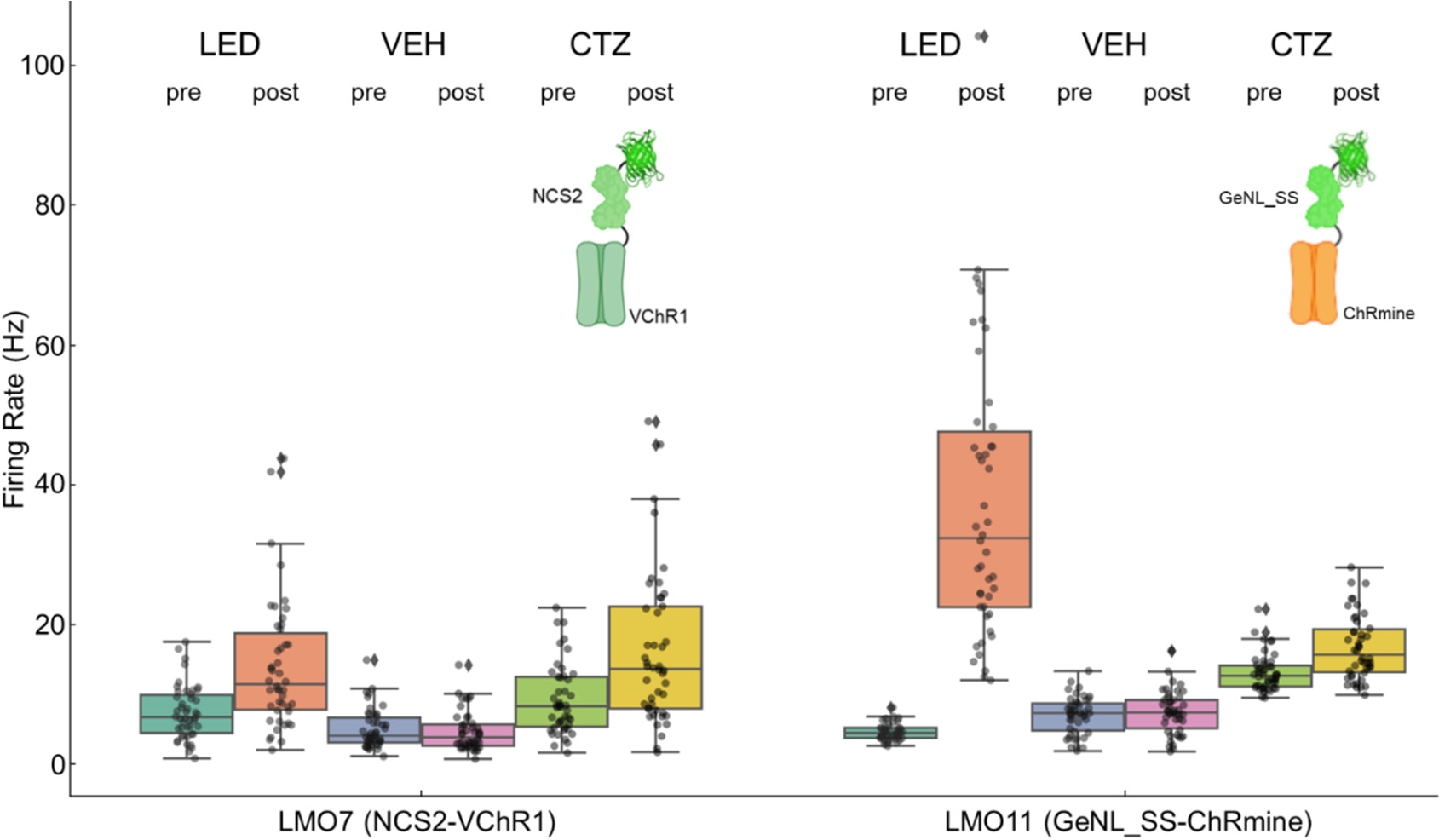
Multi-electrode recordings from cortical neurons transduced with AAV-hSyn-LMO7 and -LMO11. Average spiking frequencies of MEA channels before any treatment (pre) and after application (post) of physical or biological light, or vehicle. Spiking frequency increased when neurons were exposed to an LED light and with application of 10 μM CTZ but did not change with application of vehicle.

In the LMO7 expressing neurons, the LED treatment resulted in a significant increase in firing rates from pre-treatment (mean: 6.92, 95% CI: [5.86, 7.97]) to post-treatment (mean: 12.63, 95% CI: [10.31, 14.95]), with a corrected p-value of 0.0000. As expected, the vehicle treatment did not exhibit a significant effect (pre-treatment mean: 5.09, 95% CI: [4.27, 5.90]; post-treatment mean: 4.39, 95% CI: [3.63, 5.15], p>0.05). CTZ treatment by addition of hCTZ to the cultures for a final concentration of 10 μM increased firing rates notably (pre-treatment mean: 8.52, 95% CI: [7.26, 9.78]; post-treatment mean: 14.54, 95% CI: [11.94, 17.13], corrected p-value = 0.0002).

Similarly, in the LMO11 group, the LED treatment indicated a profound elevation in firing rates (pre-treatment mean: 4.48, 95% CI: [4.14, 4.81]; post-treatment mean: 34.75, 95% CI: [29.11, 40.39], corrected p-value of 0.0000. The vehicle treatment showed a slight increase that did not reach statistical significance (pre-treatment mean: 6.75, 95% CI: [5.88, 7.61]; post-treatment mean: 7.10, 95% CI: [6.16, 8.04], corrected p-value p>0.05). In contrast, the CTZ treatment showed a significant enhancement of neural activity (pre-treatment mean: 12.91, 95% CI: [12.20, 13.62]; post-treatment mean: 16.65, 95% CI: [15.13, 18.17], corrected p-value=0.0002).

These results indicate that the optogenetic elements in LMOs 7 and 11 can be activated by both physical and biological light sources and that bioluminescence can provide sufficient excitatory drive to increase spontaneous activity in a population of neurons.

### 3.7 LMO coupling efficiency versus bioluminescence light emission

We previously found that radiance decreased for opsin-tethered light emitters, consistent with a loss of photon emission to direct energy transfer to the nearby opsin chromophore^21^. We further observed a larger decrease for LMO7 (NCS2-VChR1) versus LMO3 (sbGLuc-VChR1). This is consistent with FRET from light emitter to opsin being more efficient for NCS2 to VChR1 (LMO7) than for sbGLuc to VChR1 (LMO3).

We saw similar tendencies with the new set of LMOs tested here. The two LMOs that basically lacked coupling efficiencies, LMO7.3 (NCS2-VChR1-NCS2) and LMO8 (NCS2-ChRger3), displayed the highest per cell bioluminescence, with LMO7.3 showing the highest light emission, probably due to the two copies of NCS2 per opsin. In contrast, LMOs with efficient coupling (LMO10, LMO11) had decreased radiance, presumably due to better FRET compatibility with the opsin chromophore (GeNL_SS > VChR1, GeNL_SS > ChRmine).

## 4 Discussion

Here we assessed the efficiencies in changing membrane potential of various LMOs consisting of different pairings of bioluminescent light emitters and light sensing opsins. Through whole cell patch clamp recordings in HEK293 cells we determined the coupling efficiency for all tested LMOs, the amplitude of bioluminescence-driven inward current compared to the maximal photocurrent elicited by a physical light source (see [Fig. 10] for summary). Through multielectrode array recordings in primary neurons we confirmed that the findings in HEK293 cells broadly translated to application in neuronal populations. By comparing LMOs that differed in specific features, such as brightness of the light emitter, sensitivity of the light sensor, arrangement of moieties, and compatibility of emission/absorption spectra, we identified guiding principles of engineering LMOs that will be useful for future LMO designs.

Brightness of the light emitter was identified as critical early on in LMO designs^5^. We and others are continuously working on generating ever brighter luminescent probes for multiple applications, including imaging with intact and split luciferases^25,31^. We focused on engineering highly efficient FRET pairs between a luciferase donor and a fluorescent protein acceptor^25^. Pairing of novel luciferase variants obtained through molecular evolution with mNeonGreen resulted in bright FRET-based light emission (sbALuc1-mNeonGreen, NCS1^21^; mNeonGreen-eKL9h, NCS2^21^; mNeonGreen-SSLuc, GeNL_SS25^25^). Interestingly, when paired with VChR1 in an LMO, the coupling efficiency of NCS2 to VChR1 (LMO7) proved to be significantly higher than for NCS1 (LMO6), suggesting that the orientation of luciferase and fluorescent protein relative to the optogenetic channel may also be critical (i.e., mNeonGreen-eKL9h-VChR1 is superior to sbALuc1-mNeonGreen-VChR1)^21^. We applied this principle successfully with LMOs 10 and 11 (mNeonGreen-SSLuc and GeNL_SS fused to VChR1 and ChRmine, respectively; this study).

It seems intuitively obvious that using a more light-sensitive opsin should increase coupling efficiency of LMOs. This has been borne out previously^3,20,22^ and again in this study (LMO 10 versus LMO11). The only exception to this rule thus far is LMO8, which uses the designer opsin ChRger3^23^. It is possible that the machine learning parameters chosen for an improved channelrhodopsin activated by a physical light source led to a reorientation of the retinal chromophore, making FRET-based activation by biological light sources less efficient.

In principle, the light emitter can be tethered to the N- or C-terminus of the opsin, and both options have led to robustly working LMOs. For instance, in our previous designs the light emitter was always tethered to the N-terminus of the opsin, but for the inhibitory iLMO2 Tung et al. fused it to the C-terminus^4^. As the arrangement of moieties was never investigated in direct comparisons, we generated and tested LMO7 variants with N- (LMO7), C- (LMO7.2), or N- and C- (LMO7.3) terminal fusions of NCS2 to VChR1. Moving the light emitter to the C-terminus in LMO7.2 caused a considerable drop in coupling efficiency compared to the N-terminal fusion (LMO7). While access to the luciferin is likely a minor factor, as luciferins cross the cell membrane efficiently, the previous observation of differences due to orientation of luciferase and fluorescent protein relative to the channel may be critical here as well (LMO7: <u>mNeonGreen-eKL9h</u>-VChR1 versus LMO7.2: VChR1-<u>mNeonGreen-eKL9h</u>). Interestingly, attaching the light emitter to both termini seemed to interfere with the proper trafficking of the channel to the membrane (see representative fluorescent image of LMO7.3-expressing HEK cell [Fig. 6(c)]) and possibly also with proper folding of the channel protein. A weak photocurrent was observed only sometimes, and bioluminescence-induced inward current could not be generated at all. In contrast, bioluminescence radiance from the two luciferases remained intact.

It appears obvious that this class of probes should benefit from higher compatibility of absorption and emission wavelengths of opsin and light emitter. However, most opsins have very wide absorbance spectra that extend activation far beyond their reported peak wavelengths, with high likelihood of activation across blue, green, and red wavelengths alike. VChR1, with an absorption peak of 535 nm, can be activated efficiently with sbGLuc (emission peak 480 nm), but activation is even more efficient with light emitters with emission peaks of 520 nm (NCS1, NCS2, GeNL_SS). Within the group of emitters with similar emission wavelength we found significant differences in coupling efficiencies (NCS1 < NCS2 < GeNL_SS; reference 21 and this study) that cannot be explained simply by matching spectra between emitter and opsin. Another example is ChRmine that has an absorption peak at 585 nm. When paired with both a green (520 nm) and a red-shifted (589 nm) light emitter, ChRmine performs best in LMO11 with the green, brightest, N-terminally tethered light emitter (GeNL_SS) rather than with the dimmer, C-terminally tethered, but most well-matched emission wavelength emitter (mCh22.0-RLuc8.6) in LMO12. Some of these discrepancies may be due to differing folding efficiencies of the light emitters tested.

Overlapping emission and absorbance spectra of the donor and an acceptor in Förster resonance energy transfer (FRET) is known to be of critical importance to maximize energy transfer efficiency. This has been studied mainly for FRET between small molecule fluorophores, between fluorescent proteins, or between a fluorescent and a bioluminescent protein. We hypothesize that FRET is the mechanism underlying the most efficient bioluminescent activation of opsins since this class of probes works reliably despite the fact that the absolute intensity of illumination by biological light sources is far lower than that which can be achieved using LEDs or lasers. If proximity of the two moieties is causing radiationless resonance energy transfer rather than simple, photon-mediated energy transfer, then the donor, the light emitter, will show loss of photon emission due to direct energy transfer to the opsin with efficient pairing, while it will recover emission when the acceptor, the opsin, is a less effective acceptor. This is supported by our observations that radiance of LMOs increases with decreasing coupling efficiency and vice versa (reference 21 and this study [Fig. 9]). This is further supported by experiments carried out by Berglund et al. 2020, where the luciferase moiety recovered bioluminescence after it was detached from the acceptor opsin through protease cleavage of the linker between luciferase and opsin^22^. Future developments of improved LMOs might benefit from molecular evolution of fusion proteins to select for the most compatible FRET pairs of light emitters and opsins.

**Fig. 9.**
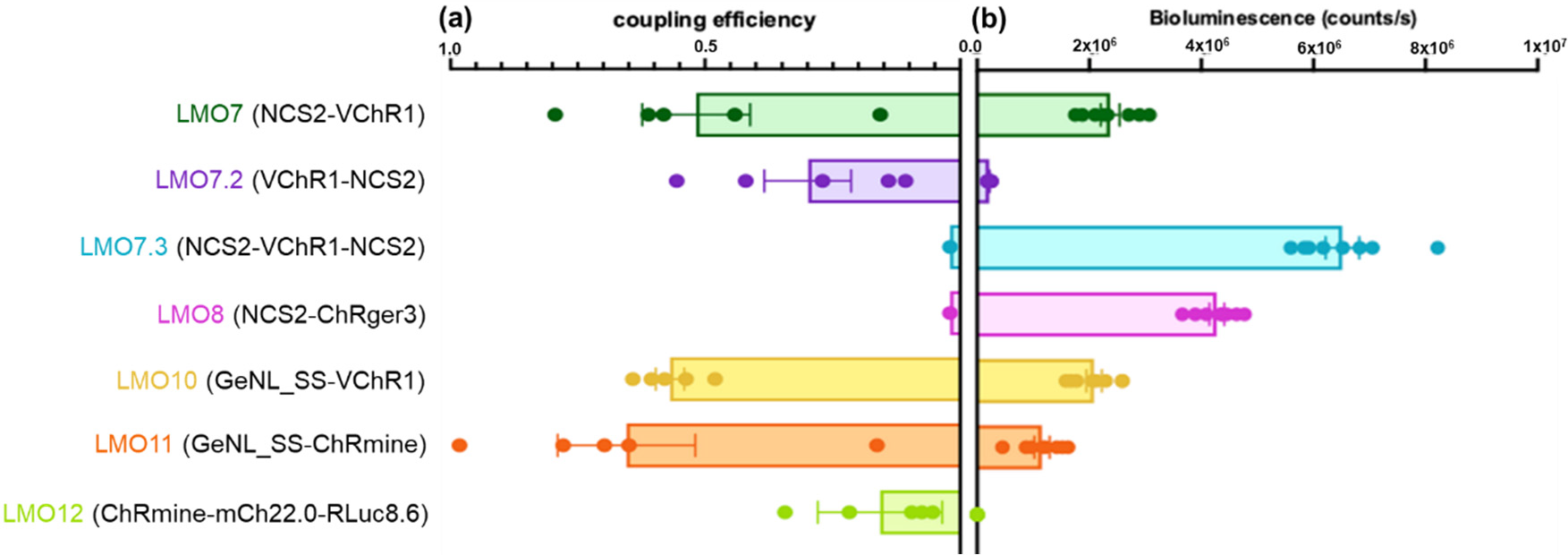
Coupling efficiency versus bioluminescence produced by LMOs. Coupling efficiencies as determined throughout the above studies are plotted on the left, and bioluminescence measured from each LMO is plotted to the right. Bioluminescence produced by each LMO was measured in HEK293 cells in a plate reader with 100 μM concentration of luciferin (n=8 for each LMO). Readings were taken immediately after addition of substrate. Bioluminescence was normalized to expression of fluorescent reporter in each LMO using fluorescence ROI analysis in ImageJ.

**Fig. 10.**
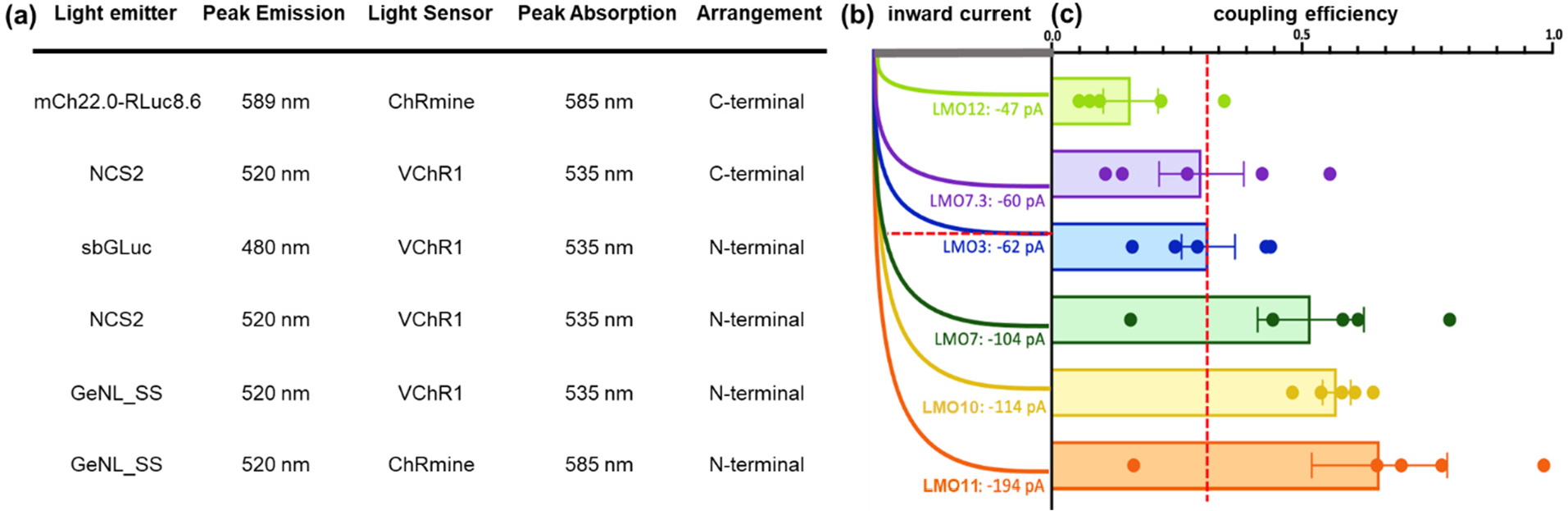
Summary of patch clamp recordings in HEK293 cells. (a) Key features of LMOs tabulated in the order displayed in (b) and (c). (b) Representative example of luciferin-induced inward currents for each LMO tested. (c) Coupling efficiency graphs of LMOs tested. Red line in (b) and (c) indicates the level of performance of LMO3, the previously determined standard for a robustly functioning LMO.

Ongoing advancements in identifying and evolving optogenetic actuators^24,32^, bioluminescent light emitters^25^, and luciferins enabling increased light emission and better penetration into the brain^33,34^ offer a range of options for improving LMOs and broadening bioluminescent optogenetics applications. Our findings provide a basis for future engineering of highly efficient LMOs that tether bright fluorescent protein – luciferase constructs N-terminally to exceedingly light-sensitive opsins, selected for optimal FRET pairing through molecular evolution of the fusion proteins. We identified LMO11 (GeNL_SS-ChRmine) as a new and improved LMO for use in bioluminescent optogenetics based projects. Lastly, we demonstrate that whole cell patch clamp recordings in HEK293 cells is an efficient strategy to quickly screen through LMO constructs.

## Data Availability Statement

The data in this manuscript will be shared in DANDI.

## Acknowledgments

We would like to thank all members of the Bioluminescence Hub (http://www.bioluminescence_hub.org/) laboratories for their feedback, discussions, and thoughtful comments throughout. This work was funded by the National Institutes of Health (R21MH101525, R01GM121944, and U01NS099709), National Science Foundation (CBET-1464686, DBI-1707352), and the W.M. Keck Foundation.

